# Performance of neural networks for prediction of asparagine content in wheat grain from imaging data

**DOI:** 10.1101/2023.12.04.569839

**Authors:** Joseph Oddy

## Abstract

**Background:** The prediction of desirable traits in wheat from imaging data is an area of growing interest thanks to the increasing accessibility of remote sensing technology. However, as the amount of data generated continues to grow, it is important that the most appropriate models are used to make sense of this information. Here, the performance of neural network models in predicting grain asparagine content is assessed against the performance of other models.

**Results:** Neural networks had greater accuracies than partial least squares regression models and gaussian naïve Bayes models for prediction of grain asparagine content, yield, genotype, and fertiliser treatment. Genotype was also more accurately predicted from seed data than from canopy data.

**Conclusion:** Using wheat canopy spectral data and combinations of wheat seed morphology and spectral data, neural networks can provide improved accuracies over other models for the prediction of agronomically important traits.

## INTRODUCTION

The global food system is highly dependent on a continuous and predictable supply of crop harvests and trade in order to operate properly. Disruptions to crop harvests (e.g. caused by losses from biotic and abiotic stressors, conflict, and export restrictions) can contribute to food insecurity, but through accurate forecasting some of these disruptions can be predicted in advance^1^. Forecasting enables actors in the food supply chain (e.g. farmers, government officials, NGOs) to put appropriate measures in place before disruption is experienced, helping to mitigate the impacts of food insecurity.

In most situations, forecasting aims to predict future crop yields as this is often the clearest metric by which to measure food insecurity^2^. However, many other metrics beyond yield are required to assess food insecurity, including measurements of food quality. One of these metrics for quality in wheat is the concentration of asparagine in grain, because asparagine is the key precursor to the processing contaminant acrylamide in wheat-based food products. Consequently, food manufacturers can minimise acrylamide formation by using raw materials (wheat flour) with low concentrations of asparagine. However, grain asparagine content is difficult to measure, so this data is normally not available.

Previous studies have found that asparagine content can be minimised through certain genetic and agronomic controls^3^ but, whilst these can be effective, these methods do not allow us to predict asparagine content. Recently, it was demonstrated that crop canopy and seed imaging technologies could be combined with modelling to enable prediction of asparagine content or (in the case of seed imaging) classification into high versus low asparagine classes^4^. The development of models to predict asparagine content in that study yielded promising results, so the continued refinement and testing of such data may help to illuminate which models can predict asparagine content most accurately.

Partial Least Squares Regression (PLSR) is often the method of choice for researchers modelling variables from spectral data^5^, however artificial neural networks are also gaining popularity for modelling spectral data. For example, Vásquez et al.^6^ showed that neural networks were more accurate than PLSR models when predicting cheese hardness from hyperspectral data. Houngbo et al.^7^ similarly found that neural networks outperformed PLSR models for the prediction of amylose content in yam from near infra-red spectroscopy (NIRS) data. Neural networks can also outperform when it comes to classification of seed quality from combinations of spectral and morphological data. This was recently illustrated by Buenafe et al.^8^, who used artificial neural network models on a morphological and spectral dataset of rice grains to improve classifications of rice quality over random forest models.

In this study, the accuracy of PLSR models and classification models (for prediction of grain asparagine content from canopy spectral data and seed morphology/spectral data, respectively) were compared with neural networks to assess which models may be most appropriate for future model development.

## MATERIALS AND METHODS

### Data collection

Data were obtained from Oddy et al.^4^; a full discussion of how those data were generated and the underlying trial structure is available in the same paper. Briefly, for the canopy spectral data, six readings were taken at intervals of approximately 2 weeks apart in 2021 from May to August using a Tec5 HandySpec Field spectrometer (Oberursel, Germany). This yielded spectral reflectance data at 10 nm intervals from 360 to 1000 nm. For the seed data, a subset of grain samples was run through a Videometer SeedLab system (Videometer, DK), yielding a mixture of morphological and spectral reflectance data. The full list of variables can be found in the corresponding datasets in the supporting information.

### Modelling

Asparagine data (mmol/kg) were loge transformed before modelling due to positive skew whilst yield data (kg/ha) were left untransformed. All models were implemented from the sci-kit learn package^9^ in python and all predictor variables were scaled before modelling using the standard scaler in sci-kit learn. The PLSR models for asparagine and yield were specified with the same number of components used in the previous study (10 for asparagine and 3 for yield). Gaussian naïve Bayes (GNB) models were similarly used for the classification tasks as done previously. For sulphur treatment classification, samples were classified into two categories: those that received 0 kg/ha of sulphur fertiliser and those that received ≥ 10 kg/ha of sulphur fertiliser.

Neural networks were implemented as multi-layer perceptrons (MLPs). To help guide selection of appropriate MLP structures, grid search cross-validation was performed for each MLP to find the best parameters out of a specified parameter space. Further details of mparameter spaces used for model selection are available in Jupyter Notebook files in the supporting information. Details of MLP structures chosen following grid search are provided in Table 1. Model performance metrics (*R*^2^ for regression models and balanced accuracy for classification models) were calculated by five-fold cross validation repeated 100 times. Data manipulation and plotting was carried out using NumPy^10^, pandas^11^, and plotnine.

**Table 1.** Details of models used in this study and key model performance metrics (as means). PLSR (partial least squares regression), MLP (multi-layer perceptron), GNB (gaussian naïve Bayes). *R*^2^ is given for regression models whilst balanced accuracy metrics are given for classification models.

| Data | Target variable | Model | Parameters | $R^2$/Accuracy |
| --- | --- | --- | --- | --- |
| Canopy | Asparagine content | PLSR | 10 components | 71.3% |
| | | MLP Regressor | 2-layers × 300 neurons | 74.8% |
|  | Yield | PLSR | 3 components | 79.8% |
| | | MLP Regressor | 3-layers × 50 neurons | 84.5% |
|  | Sulphur treatment | GNB | None | 75.9% |
| | | MLP Classifier | 1-layer × 100 neurons | 87.5% |
|  | Genotype | GNB | None | 26.5% |
| | | MLP Classifier | 1-layer × 300 neurons | 31.7% |
| Seed | Sulphur treatment | GNB | None | 85.9% |
| | | MLP Classifier | 1-layer × 50 neurons | 88.9% |
|  | Genotype | GNB | None | 40.4% |
| | | MLP Classifier | 2-layers × 50 neurons | 74.0% |

## RESULTS

Firstly, the accuracy of MLP models for predicting asparagine and yield from spectral canopy data was tested relative to PLSR models. For the prediction of asparagine content, the neural network was more accurate by approximately 3.7% on average, whilst for prediction of yield, the neural network was more accurate by approximately 4.7% (Table 1). Results from the five-fold cross validation are also shown in Figure 1a.

**Figure 1.**
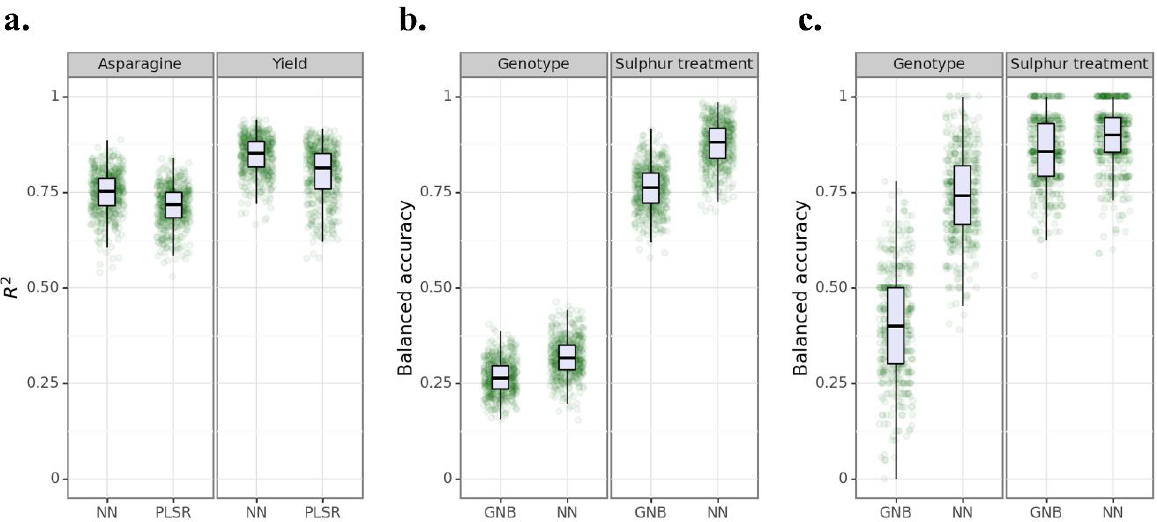
Performance of neural networks (NNs) relative to other models used for prediction from imaging data as illustrated by five-fold cross validation accuracy scores. **a**. *R*^2^ of models predicting asparagine content (mmol per kg) and yield (tonnes per ha) from canopy spectral data. **b**. Balanced accuracies of models predicting genotype and sulphur treatment categories from canopy spectral data. **c**. Balanced accuracies of models predicting genotype and sulphur treatment categories from seed morphology/spectral data. PLSR (partial least squares regression), GNB (gaussian naïve Bayes).

Secondly, the accuracy of neural networks for classification of samples by genotype and sulphur treatment was assessed using the canopy spectral data. Again, neural networks were more accurate than GNB models for prediction of genotype and sulphur treatment class, with mean improvements in accuracy of 5.2% and 11.6%, respectively (Table 1; Figure 1b). The accuracy of models for genotype prediction were much lower than those for sulphur treatment prediction, likely due to the number of categories required for classification in each scenario (12 genotypes for classification relative to only 2 sulphur treatment categories).

Finally, the accuracy of neural networks for classification of samples by genotype and sulphur treatment was assessed using the seed spectral/morphological data. Here, neural networks also had greater accuracies than GNB models, with mean improvements of 3% for sulphur treatment classification and, notably, 33.6% for genotype classification (Table 1; Figure 1c). In general, classification models using the seed data were more accurate than the classification models built using canopy data.

## DISCUSSION

Beyond the data used in this study, only one other study has previously attempted to predict grain asparagine content from imaging data. Rapp et al.^12^ used NIRS in their study, however they found that their model was unable to accurately predict grain asparagine content. In contrast, the PLSR model presented previously^4^ and the neural network presented here were able to predict grain asparagine content with accuracies of 71.3% and 74.8%, respectively. Consequently, collection of spectral data and model building should be continued in order to further improve these predictive models and to enable their use in commercial farming and food manufacturing. For example, development of models that are able to detect potentially high asparagine forming grains early enough in the growing season could allow farmers to implement strategies to reduce asparagine formation (e.g. through application of fertilisers and disease control). Similarly, development of models for predicting the asparagine content of seeds would allow food manufacturers and millers to use only those grains that are lower in asparagine content, thereby minimising acrylamide formation in their products.

More broadly, the results of this analysis are consistent with previous studies showing that neural network models can outperform other models for the prediction of agricultural and food quality data^6-8^. The greatest improvement in this study was observed between GNB and neural network models for the prediction of genotype from seed data. In most other cases, neural networks outperformed the PLSR or GNB models by under 10%, however in this scenario the neural network outperformed the GNB model by 33.6%, boosting the predictive accuracy from 40.4% to 74.0%. The neural network model trained on the canopy data was not able to make similarly large increases in prediction of genotype over the GNB model, indicating that seed data is better suited for distinguishing genotypes than canopy measurements. This differs from the findings of Gao et al.^13^, who found that images of wheat during tillering and flowering enabled more accurate classification of wheat genotypes than images of seeds. However, that study also used RGB images as input instead of decomposed spectral/morphological data (as is used in this study), which may account for some of these differences.

## CONCLUSIONS

Overall, the neural network models used in this study were able to more accurately predict wheat grain asparagine content, yield, genotype, and fertiliser treatment than either PLSR or GNB models. Consequently, researchers should consider using neural networks when developing tools for predicting traits from wheat canopy and seed imaging data.

## ACKNOWLEDGEMENTS

I am grateful for the guidance and advice of Nigel Halford, under whose guidance the data used in this study were originally generated, and for scholarship support from the Society of Chemical Industry.

