## Supplementary material for "Performance of neural networks for prediction of asparagine content in wheat grain from imaging data": 230704 NN seed data.ipynb

```
# %% [markdown]
# DATA IMPORT AND PRE-PROCESSING

# %% [code]
#%% import required packages
import numpy as np
import pandas as pd
from sklearn import model_selection
from sklearn.model_selection import RepeatedKFold
from sklearn.naive_bayes import GaussianNB
from sklearn.neighbors import KNeighborsClassifier
import os
from plotnine import ggplot, aes, geom_boxplot, geom_jitter, theme_bw, theme, xlab, ylab, facet_wrap, ylim
from sklearn.neural_network import MLPClassifier
from sklearn.model_selection import GridSearchCV
from sklearn.model_selection import cross_validate

# %% [code]
# Import required libraries
import pandas as pd
import numpy as np 
import matplotlib.pyplot as plt
import sklearn
from sklearn.neural_network import MLPClassifier
from sklearn.neural_network import MLPRegressor

# Import necessary modules
from sklearn.model_selection import train_test_split
from sklearn.metrics import mean_squared_error
from math import sqrt
from sklearn.metrics import r2_score

# %% [code]
#%% set directory
os.getcwd()
os.chdir('C:\\Users\\BIOL0050\\OneDrive - Nexus365\\0.7. Data sci\\MDPI Crops analysis\\')

# %% [code]
#%% import data
data_full = pd.read_csv('220907 all data.csv')
data_full = pd.DataFrame(data_full)
data = data_full.iloc[:, 5:408]

# %% [code]
from sklearn.preprocessing import StandardScaler
scaler = StandardScaler() 

#%% define predictor and response variables
X = data.iloc[:, 1:405].values
X = scaler.fit_transform(X)
y = data_full[["nutrient_treatment"]].values.ravel()
yvar = data_full[["cultivar"]].values.ravel()

# %% [markdown]
# SULPHUR PREDICTION

# %% [code]
# set up neural network model and investigate optimal parameters (for agronomic treatment prediction)
mlp = MLPClassifier(max_iter=1000)
parameter_space = {
    'hidden_layer_sizes': [(50), (100), (50,50), (100,100), (50, 50, 50), (100, 100, 100)],
    'activation': ['tanh', 'relu'],
    'solver': ['sgd', 'adam'],
    'alpha': [0.0001, 0.05, 1],
    'learning_rate': ['constant','adaptive'],
}

clf = GridSearchCV(mlp, parameter_space, n_jobs=-1, cv=5)
clf.fit(X, y)

# Best parameter set
print('Best parameters found:\n', clf.best_params_)

# %% [code]
#%% model for predicting agronomic treatment
#define cross-validation method
cv = RepeatedKFold(n_splits = 5, n_repeats = 100)
# define models
model = GaussianNB()
model2 = MLPClassifier(activation='tanh', alpha=0.05, hidden_layer_sizes=(50), learning_rate = 'constant', solver='sgd')

# %% [code]
# evaluate models
scoring = {'bal_acc':'balanced_accuracy'}
scores = cross_validate(model, X, y, scoring=scoring, cv=cv, n_jobs = -1)
scores2 = cross_validate(model2, X, y, scoring=scoring, cv=cv, n_jobs = -1)

# %% [code]
# summarize result
print(np.mean(scores['test_bal_acc']))
print(np.mean(scores2['test_bal_acc']))

# %% [code]
#%% extract and combine model accuracy scores
gnbdf = pd.DataFrame(scores['test_bal_acc'])
gnbdf.columns = ['GNB']
nndf = pd.DataFrame(scores2['test_bal_acc'])
nndf.columns = ['NN']
fulldata = pd.concat([gnbdf, nndf], axis=1)

# %% [code]
print(fulldata)

# %% [code]
#%% reshape data
fulldata2 = fulldata.melt(var_name = 'model', value_name = 'Accuracy')

# %% [code]
#%% plot accuracy scores
(ggplot(fulldata2, aes(x = 'model', y = 'Accuracy'))
 + geom_jitter(width = 0.25, alpha = 0.05, color = "darkgreen")
 + geom_boxplot(width = 0.25, color = "black", fill = "lavender", outlier_shape = 'x', outlier_colour = '') 
 + theme_bw()
 + theme(figure_size=(2, 4))
 + xlab("")
 + ylab("Balanced accuracy")
 )

# %% [markdown]
# CULTIVAR PREDICTION

# %% [code]
# set up neural network model and investigate optimal parameters (for cultivar prediction)
mlp = MLPClassifier(max_iter=1000)
parameter_space = {
    'hidden_layer_sizes': [(50), (100), (50,50), (100,100), (50, 50, 50), (100, 100, 100)],
    'activation': ['tanh', 'relu'],
    'solver': ['sgd', 'adam'],
    'alpha': [0.0001, 0.05, 1],
    'learning_rate': ['constant','adaptive'],
}

clf = GridSearchCV(mlp, parameter_space, n_jobs=-1, cv=5)
clf.fit(X, yvar)

# Best parameter set
print('Best parameters found:\n', clf.best_params_)

# %% [code]
#%% models for predicting cultivar
#define cross-validation method
cv = RepeatedKFold(n_splits = 5, n_repeats = 100)
# define models
varmodel = GaussianNB()
varmodel2 = MLPClassifier(activation='tanh', alpha=1, hidden_layer_sizes=(50,50), learning_rate = 'constant', solver='adam')
varmodel3 = KNeighborsClassifier()

# %% [code]
# evaluate models
scoring = {'bal_acc':'balanced_accuracy'}
varscores = cross_validate(varmodel, X, yvar, scoring=scoring, cv=cv, n_jobs = -1)
varscores2 = cross_validate(varmodel2, X, yvar, scoring=scoring, cv=cv, n_jobs = -1)

# %% [code]
# summarize result
print(np.mean(varscores['test_bal_acc']))
print(np.mean(varscores2['test_bal_acc']))

# %% [code]
#%% extract and combine model accuracy scores
vargnbdf = pd.DataFrame(varscores['test_bal_acc'])
vargnbdf.columns = ['GNB']
varnndf = pd.DataFrame(varscores2['test_bal_acc'])
varnndf.columns = ['NN']
varfulldata = pd.concat([vargnbdf, varnndf], axis=1)

# %% [code]
#%% reshape data
varfulldata2 = varfulldata.melt(var_name = 'model', value_name = 'Accuracy')

# %% [code]
#%% plot accuracy scores
(ggplot(varfulldata2, aes(x = 'model', y = 'Accuracy'))
 + geom_jitter(width = 0.25, alpha = 0.05, color = "darkgreen")
 + geom_boxplot(width = 0.25, color = "black", fill = "lavender", outlier_shape = 'x', outlier_colour = '') 
 + theme_bw()
 + theme(figure_size=(2, 4))
 + xlab("")
 + ylab("Balanced accuracy")
 )

# %% [markdown]
# PLOTTING TOGETHER

# %% [code]
fulldata2['variable'] = "Sulphur treatment"
varfulldata2['variable'] = "Genotype"
completedata = pd.concat([fulldata2, varfulldata2])

# %% [code]
print(completedata)

# %% [code]
(ggplot(completedata, aes(x = 'model', y = 'Accuracy'))
 + geom_jitter(width = 0.25, alpha = 0.05, color = "darkgreen")
 + geom_boxplot(width = 0.25, color = "black", fill = "lavender", outlier_shape = 'x', outlier_colour = '') 
 + theme_bw()
 + facet_wrap('variable')
 + theme(figure_size=(3, 4))
 + xlab("")
 + ylab("Balanced accuracy")
 + ylim(0, 1)
 )

# %% [code]
```
