## Supplementary material for "Performance of neural networks for prediction of asparagine content in wheat grain from imaging data": 230728 NN spectral data.ipynb

```
# %% [markdown]
# DATA IMPORT

# %% [code]
#%% import packages
import numpy as np
import pandas as pd
import matplotlib.pyplot as plt
from sklearn import model_selection
from sklearn.model_selection import RepeatedKFold
from sklearn.cross_decomposition import PLSRegression
from sklearn.naive_bayes import GaussianNB
import os
from plotnine import ggplot, aes, geom_boxplot, theme_bw, theme, geom_violin, xlab, ylab, ylim
from plotnine import ggplot, aes, geom_boxplot, geom_jitter, theme_bw, theme, xlab, ylab, facet_wrap
from sklearn.preprocessing import StandardScaler
scaler = StandardScaler() 
from sklearn.neural_network import MLPRegressor
from sklearn.neural_network import MLPClassifier
from sklearn.model_selection import GridSearchCV
from sklearn.model_selection import cross_validate

# %% [code]
#%% set directory
os.getcwd()
os.chdir('C:\\Users\\BIOL0050\\OneDrive - Nexus365\\0.7. Data sci\\MDPI Crops analysis')

# %% [code]
#%% import data
data_full = pd.read_csv('220721 all spectral readings.csv')
data_full = pd.DataFrame(data_full)
data = data_full.iloc[:, 4:408]
data = data[data['ln.asn.average'].notna()]

# %% [code]
#%% define predictor and response variables for asn and yield
X = data.iloc[:, 2:405].values
X = scaler.fit_transform(X)
y = data[["ln.asn.average"]].values.ravel()
yyield = data[["Grain85%"]].values.ravel()

# %% [markdown]
## ASPARAGINE PREDICTION

# %% [code]
# set up neural network model and investigate optimal parameters (for agronomic treatment prediction)
mlp = MLPRegressor(max_iter=1000)
parameter_space = {
    'hidden_layer_sizes': [(50), (50,50), (50,50,50), (100), (100,100), (100,100,100), (300), (300,300), (300,300,300)],
    'activation': ['tanh', 'relu'],
    'solver': ['sgd', 'adam'],
    'alpha': [0.0001, 0.05, 1],
    'learning_rate': ['constant','adaptive'],
}

clf = GridSearchCV(mlp, parameter_space, n_jobs=-1, cv=5)
clf.fit(X, y)

# Best parameter set
print('Best parameters found:\n', clf.best_params_)

# %% [code]
#%% model for predicting agronomic treatment
#define cross-validation method
cv = RepeatedKFold(n_splits = 5, n_repeats = 100)
# define models
model = PLSRegression(n_components = 10)
model2 = MLPRegressor(activation='relu', alpha=1, hidden_layer_sizes=(300, 300), learning_rate = 'adaptive', solver='adam')

# %% [code]
# evaluate models
scoring = {'r2':'r2', 'RMSE':'neg_root_mean_squared_error', 'explained_variance': 'explained_variance'}
scores = cross_validate(model, X, y, scoring=scoring, cv=cv, n_jobs = -1)
scores2 = cross_validate(model2, X, y, scoring=scoring, cv=cv, n_jobs = -1)

# %% [code]
# summarize result
print(scores.keys())
print(np.mean(scores['test_r2']))
print(np.mean(scores2['test_r2']))
print(np.mean(scores['test_RMSE']))
print(np.mean(scores2['test_RMSE']))
print(np.mean(scores['test_explained_variance']))
print(np.mean(scores2['test_explained_variance']))

# %% [code]
#%% extract and combine model accuracy scores
plsrdf = pd.DataFrame(scores['test_r2'])
plsrdf.columns = ['PLSR']
nndf = pd.DataFrame(scores2['test_r2'])
nndf.columns = ['NN']
fulldata = pd.concat([plsrdf, nndf], axis=1)

# %% [code]
print(fulldata)

# %% [code]
#%% reshape data
fulldata2 = fulldata.melt(var_name = 'model', value_name = 'Accuracy')

# %% [code]
#%% plot accuracy scores
(ggplot(fulldata2, aes(x = 'model', y = 'Accuracy'))
 + geom_jitter(width = 0.25, alpha = 0.05, color = "darkgreen")
 + geom_boxplot(width = 0.25, color = "black", fill = "lavender", outlier_shape = 'x', outlier_colour = '') 
 + theme_bw()
 + theme(figure_size=(2, 4))
 + xlab("")
 + ylab("R2")
 )

# %% [markdown]
## YIELD PREDICTION

# %% [code]
# set up neural network model and investigate optimal parameters (for agronomic treatment prediction)
mlp = MLPRegressor(max_iter=1000)
parameter_space = {
    'hidden_layer_sizes': [(50), (50,50), (50,50,50), (100), (100,100), (100,100,100), (300), (300,300), (300,300,300)],
    'activation': ['tanh', 'relu'],
    'solver': ['sgd', 'adam'],
    'alpha': [0.0001, 0.05, 1, 10],
    'learning_rate': ['constant','adaptive'],
}

clf = GridSearchCV(mlp, parameter_space, n_jobs=-1, cv=5)
clf.fit(X, yyield)

# Best parameter set
print('Best parameters found:\n', clf.best_params_)

# %% [code]
#%% model for predicting agronomic treatment
#define cross-validation method
cv = RepeatedKFold(n_splits = 5, n_repeats = 100)
# define models
yieldmodel = PLSRegression(n_components = 3)
yieldmodel2 = MLPRegressor(activation='tanh', alpha=10, hidden_layer_sizes=(50,50,50), learning_rate = 'constant', solver='adam')

# %% [code]
# evaluate models
scoring = {'r2':'r2', 'RMSE':'neg_root_mean_squared_error', 'explained_variance': 'explained_variance'}
yieldscores = cross_validate(yieldmodel, X, yyield, scoring=scoring, cv=cv, n_jobs = -1)
yieldscores2 = cross_validate(yieldmodel2, X, yyield, scoring=scoring, cv=cv, n_jobs = -1)

# %% [code]
# summarize result
print(scores.keys())
print(np.mean(yieldscores['test_r2']))
print(np.mean(yieldscores2['test_r2']))

# %% [code]
#%% extract and combine model accuracy scores
yieldplsrdf = pd.DataFrame(yieldscores['test_r2'])
yieldplsrdf.columns = ['PLSR']
yieldnndf = pd.DataFrame(yieldscores2['test_r2'])
yieldnndf.columns = ['NN']
yieldfulldata = pd.concat([yieldplsrdf, yieldnndf], axis=1)

# %% [code]
#%% reshape data
yieldfulldata2 = yieldfulldata.melt(var_name = 'model', value_name = 'Accuracy')

# %% [code]
#%% plot accuracy scores
(ggplot(yieldfulldata2, aes(x = 'model', y = 'Accuracy'))
 + geom_jitter(width = 0.25, alpha = 0.05, color = "darkgreen")
 + geom_boxplot(width = 0.25, color = "black", fill = "lavender", outlier_shape = 'x', outlier_colour = '') 
 + theme_bw()
 + theme(figure_size=(2, 4))
 + xlab("")
 + ylab("R2")
 )

# %% [markdown]
# CLASSIFICATION DATA IMPORT

# %% [code]
#read in data
data_full = pd.read_csv('220909 all spectral readings v2.csv')
data_full = pd.DataFrame(data_full)
data_full = data_full[data_full['ln.asn.average'].notna()]
data = data_full.iloc[:, 5:409]
data = data[data['ln.asn.average'].notna()]

# %% [code]
#define predictor and response variables for asn and yield
X = data.iloc[:, 2:405]
y = data_full[["binary_sulphur"]].values.ravel()
y2 = data_full[["cultivar"]].values.ravel()

# %% [markdown]
# TREATMENT PREDICTION

# %% [code]
# set up neural network model and investigate optimal parameters (for agronomic treatment prediction)
mlp = MLPClassifier(max_iter=1000)
parameter_space = {
    'hidden_layer_sizes': [(50), (50,50), (50,50,50), (100), (100,100), (100,100,100), (300), (300,300), (300,300,300)],
    'activation': ['tanh', 'relu'],
    'solver': ['sgd', 'adam'],
    'alpha': [0.0001, 0.05, 1, 10],
    'learning_rate': ['constant','adaptive'],
}


clf = GridSearchCV(mlp, parameter_space, n_jobs = -1, cv = 5, verbose = 3)
clf.fit(X, y)

# Best parameter set
print('Best parameters found:\n', clf.best_params_)

# %% [code]
#%% model for predicting agronomic treatment
#define cross-validation method
cv = RepeatedKFold(n_splits = 5, n_repeats = 100)
# define models
fertmodel = GaussianNB()
fertmodel2 = MLPClassifier(activation='tanh', alpha=0.0001, hidden_layer_sizes=(100), learning_rate = 'adaptive', solver='adam')

# %% [code]
# evaluate models
scoring = {'bal_acc':'balanced_accuracy'}
fertscores = cross_validate(fertmodel, X, y, scoring=scoring, cv=cv, n_jobs = -1)
fertscores2 = cross_validate(fertmodel2, X, y, scoring=scoring, cv=cv, n_jobs = -1)

# %% [code]
# summarize result
print(scores.keys())
print(np.mean(fertscores['test_bal_acc']))
print(np.mean(fertscores2['test_bal_acc']))

# %% [code]
#%% extract and combine model accuracy scores
fertgnbdf = pd.DataFrame(fertscores['test_bal_acc'])
fertgnbdf.columns = ['GNB']
fertnndf = pd.DataFrame(fertscores2['test_bal_acc'])
fertnndf.columns = ['NN']
fertfulldata = pd.concat([fertgnbdf, fertnndf], axis=1)

# %% [code]
print(fertfulldata)

# %% [code]
#%% reshape data
fertfulldata2 = fertfulldata.melt(var_name = 'model', value_name = 'Accuracy')

# %% [code]
#%% plot accuracy scores
(ggplot(fertfulldata2, aes(x = 'model', y = 'Accuracy'))
 + geom_jitter(width = 0.25, alpha = 0.05, color = "darkgreen")
 + geom_boxplot(width = 0.25, color = "black", fill = "lavender", outlier_shape = 'x', outlier_colour = '') 
 + theme_bw()
 + theme(figure_size=(2, 4))
 + xlab("")
 + ylab("Balanced accuracy")
 )

# %% [markdown]
# CULTIVAR PREDICTION

# %% [code]
# set up neural network model and investigate optimal parameters (for agronomic treatment prediction)
mlp = MLPClassifier(max_iter=1000)
parameter_space = {
    'hidden_layer_sizes': [(50), (50,50), (50,50,50), (100), (100,100), (100,100,100), (300), (300,300), (300,300,300)],
    'activation': ['tanh', 'relu'],
    'solver': ['sgd', 'adam'],
    'alpha': [0.0001, 0.05, 1, 10],
    'learning_rate': ['constant','adaptive'],
}

clf = GridSearchCV(mlp, parameter_space, n_jobs=-1, cv=5)
clf.fit(X, y2)

# Best parameter set
print('Best parameters found:\n', clf.best_params_)

# %% [code]
#%% model for predicting agronomic treatment
#define cross-validation method
cv = RepeatedKFold(n_splits = 5, n_repeats = 100)
# define models
genomodel = GaussianNB()
genomodel2 = MLPClassifier(activation='relu', alpha=0.0001, hidden_layer_sizes=(300), learning_rate = 'constant', solver='adam')

# %% [code]
# evaluate models
scoring = {'bal_acc':'balanced_accuracy'}
genoscores = cross_validate(genomodel, X, y2, scoring=scoring, cv=cv, n_jobs = -1)
genoscores2 = cross_validate(genomodel2, X, y2, scoring=scoring, cv=cv, n_jobs = -1)

# %% [code]
# summarize result
print(np.mean(genoscores['test_bal_acc']))
print(np.mean(genoscores2['test_bal_acc']))

# %% [code]
#%% extract and combine model accuracy scores
genognbdf = pd.DataFrame(genoscores['test_bal_acc'])
genognbdf.columns = ['GNB']
genonndf = pd.DataFrame(genoscores2['test_bal_acc'])
genonndf.columns = ['NN']
genofulldata = pd.concat([genognbdf, genonndf], axis=1)

# %% [code]
print(genofulldata)

# %% [code]
#%% reshape data
genofulldata2 = genofulldata.melt(var_name = 'model', value_name = 'Accuracy')

# %% [code]
#%% plot accuracy scores
(ggplot(genofulldata2, aes(x = 'model', y = 'Accuracy'))
 + geom_jitter(width = 0.25, alpha = 0.05, color = "darkgreen")
 + geom_boxplot(width = 0.25, color = "black", fill = "lavender", outlier_shape = 'x', outlier_colour = '') 
 + theme_bw()
 + theme(figure_size=(2, 4))
 + xlab("")
 + ylab("Balanced accuracy")
 )

# %% [markdown]
## PLOTTING TOGETHER

# %% [code]
fulldata2['variable'] = "Asparagine"
yieldfulldata2['variable'] = "Yield"
completedata = pd.concat([fulldata2, yieldfulldata2])

# %% [code]
(ggplot(completedata, aes(x = 'model', y = 'Accuracy'))
 + geom_jitter(width = 0.25, alpha = 0.05, color = "darkgreen")
 + geom_boxplot(width = 0.25, color = "black", fill = "lavender", outlier_shape = 'x', outlier_colour = '') 
 + theme_bw()
 + facet_wrap('variable')
 + theme(figure_size=(3, 4))
 + xlab("")
 + ylab("$R^2$")
 + ylim(0, 1)
 )

# %% [code]
fertfulldata2['variable'] = "Sulphur treatment"
genofulldata2['variable'] = "Genotype"
completedata2 = pd.concat([fertfulldata2, genofulldata2])

# %% [code]
(ggplot(completedata2, aes(x = 'model', y = 'Accuracy'))
 + geom_jitter(width = 0.25, alpha = 0.05, color = "darkgreen")
 + geom_boxplot(width = 0.25, color = "black", fill = "lavender", outlier_shape = 'x', outlier_colour = '') 
 + theme_bw()
 + facet_wrap('variable')
 + theme(figure_size=(3, 4))
 + xlab("")
 + ylab("Balanced accuracy")
 + ylim(0, 1)
 )

# %% [code]
```
